# Inference of Transcriptional Regulation From STARR-seq Data

**DOI:** 10.1101/2024.03.06.583826

**Authors:** Amin Safaeesirat, Hoda Taeb, Emirhan Tekoglu, Tunc Morova, Nathan A. Lack, Eldon Emberly

## Abstract

One of the primary regulatory processes in cells is transcription, during which RNA polymerase II (Pol-II) transcribes DNA into RNA. The binding of Pol-II to its site is regulated through interactions with transcription factors (TFs) that bind to DNA at enhancer cis-regulatory elements. Measuring the enhancer activity of large libraries of distinct DNA sequences is now possible using Massively Parallel Reporter Assays (MPRAs), and computational methods have been developed to identify the dominant statistical patterns of TF binding within these large datasets. Such methods are global in their approach and may overlook important regulatory sites which function only within the local context. Here we introduce a method for inferring functional regulatory sites (their number, location and width) within an enhancer sequence based on measurements of its transcriptional activity from an MPRA method such as STARR-seq. The model is based on a mean-field thermodynamic description of Pol-II binding that includes interactions with bound TFs. Our method applied to simulated STARR-seq data for a variety of enhancer architectures shows how data quality impacts the inference and also how it can find local regulatory sites that may be missed in a global approach. We also apply the method to recently measured STARR-seq data on androgen receptor (AR) bound sequences, a TF that plays an important role in the regulation of prostate cancer. The method identifies key regulatory sites within these sequences which are found to overlap with binding sites of known co-regulators of AR.

**Author Summary:** We present an inference method for identifying regulatory sites within a putative DNA enhancer sequence, given only the measured transcriptional output of a set of overlapping sequences using an assay like STARR-seq. It is based on a mean-field thermodynamic model that calculates the binding probability of Pol-II to its promoter and includes interactions with sites in the DNA sequence of interest. By maximizing the likelihood of the data given the model, we can infer the number of regulatory sites, their locations, and their widths. Since it is a local model, it can in principle find regulatory sites that are important within a local context that may get missed in a global fit. We test our method on simulated data of simple enhancer architectures and show that it is able to find only the functional sites. We also apply our method to experimental STARR-seq data from 36 androgen receptor bound DNA sequences from a prostate cancer cell line. The inferred regulatory sites overlap known important regulatory motifs and their ChIP-seq data in these regions. Our method shows potential at identifying locally important functional regulatory sites within an enhancer given only its measured transcriptional output.

## 2 Introduction

Transcription is a key process within a cell, converting genetic information in the genome into mRNA that is translated into protein. The transcription of mRNA is done by RNA polymerase II (Pol-II), a molecular motor, that is subject to regulation by a variety of mechanisms, including protein signaling, alterations in DNA structure, epigenetic modifications, and the binding of transcription factors (TFs) [1]. Notably, TFs play a crucial role in this regulatory process by binding to specific regulatory sites on the DNA and interacting with Pol-II, either directly or indirectly. These interactions are important for controlling the binding of Pol-II to its promoter site, which influences the rate of mRNA production. Identifying regulatory sites, their corresponding TFs, and how their interactions with Pol-II control transcription is an active and ongoing area of research.

Significant experimental advancements have made it possible to analyze transcription and gene regulation quantitatively on a genome-wide scale. Chromatin Immunoprecipitation Sequencing (ChIP-seq) [2] has been pivotal in identifying binding sites of TFs. Complementing this, DNase I Hypersensitive Sites Sequencing (DNase-seq) [3] and Assay for Transposase-Accessible Chromatin using sequencing (ATAC-seq) [4] have provided detailed maps of chromatin accessibility, highlighting regions likely to be involved in gene regulation. The advent of RNA Sequencing (RNA-seq) revolutionized transcriptomics by allowing for a comprehensive quantification of gene expression. The above methods can identify where TFs are bound, which sequences may be putative enhancers and how they may affect gene transcription. More recently, massively parallel reporter assays (MPRA) such as STARR-seq (Self-Transcribing Active Regulatory Region sequencing) [5, 6] have been able to quantify the enhancer activity of hundreds of thousands to millions of distinct DNA sequences in living cells. STARR-seq measures the self-transcription of putative enhancer sequences by RNA-seq, making it a quantitative assay for measuring the binding activity of Pol-II. This plasmid-based method provides a rich dataset to apply models of transcription as similar to promoter bashing assays, the varying overlap of cloned fragments allows the systematic dissection of potential regulatory elements.

Recent advances in MPRA and STARR-seq analysis have largely been driven by two complimentary methodologies. The first employs deep learning techniques [7–14], where neural networks process DNA sequences to predict the STARR-seq output [15]. These methods are able to identify known and novel binding motifs that play a part in the transcriptional program being studied, though interpretibability still remains a challenge. The second method fits a specific biophysical model [16–18] to the observed data [19, 20], often directly incorporating TF binding information. These methods provide a model which is more readily interpretable, but may miss higher order interactions or novel motifs that the neural network approach may capture. Both methodologies are categorized as global fitting approaches, finding a transcriptional regulatory model that captures architectures and grammars that are over represented in the data.

Here, we introduce an inference scheme to complement the global approaches described above, that fits a biophysical model of transcription to identify regulatory sequence elements in a putative enhancer, utilizing STARR-seq data specific to that enhancer. Given just the measured number of transcribed mRNA of a set of sequences that overlap the enhancer region of interest, our method infers the locations of regulatory sites and their effective interactions with Pol-II by maximizing the likelihood of the transcriptional data. Because it is a local method, it can potentially identify local regulatory sites that might be missed in a global fit. For instance, our model can discern scenarios where a known TF binding site might not be active in a particular enhancer, or where the effective interaction with Pol-II may depart locally from how the TF typically acts globally. This ability to capture such specific variations highlights the distinctiveness of our approach. We first apply the method to simulated STARR-seq data for a set of known enhancer architectures, made up of defined regulatory sites, to evaluate its sensitivity to the amount of data and sequencing depth. Then, we apply the inference method to experimental STARR-seq data for androgen receptor (AR) bound sequences measured in a prostate cancer cell line. We show that our method is able to identify functional regulatory sites within each of these sequences, and that they overlap well with known motifs and binding data for AR and other AR-associated TFs. Thus our de novo inference method, can readily discover regulatory sites and their effective role in transcription using only measured transcriptional output from a set of overlapping DNA sequences.

## 3 Materials and Methods

### 3.1 Biophysical model for transcription

We describe an enhancer sequence as being comprised of a set of binding sites for transcription factors (TFs) along with a binding site for RNA-Polymerase II (Pol-II). Each binding site is described by a binary variable *σ*_*i*_, that takes value *σ*_*i*_ = 1 if the site is bound, or *σ*_*i*_ = 0 if it is unbound. Additionally, each site has an on-site energy *h*_*i*_ that includes both the binding energy as well as chemical potential for the given factor. In this model, the *i*-th site can interact with the *j*-th site, with an interaction energy *J*_*i, j*_ (see Fig. 1(A)). Thus the Hamiltonian for our system is equivalent to the Sherrington-Kirkpatrick model from spin glasses and is given by

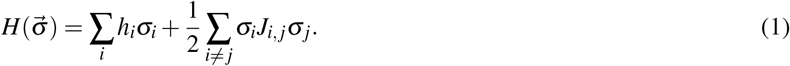

**Figure 1.**
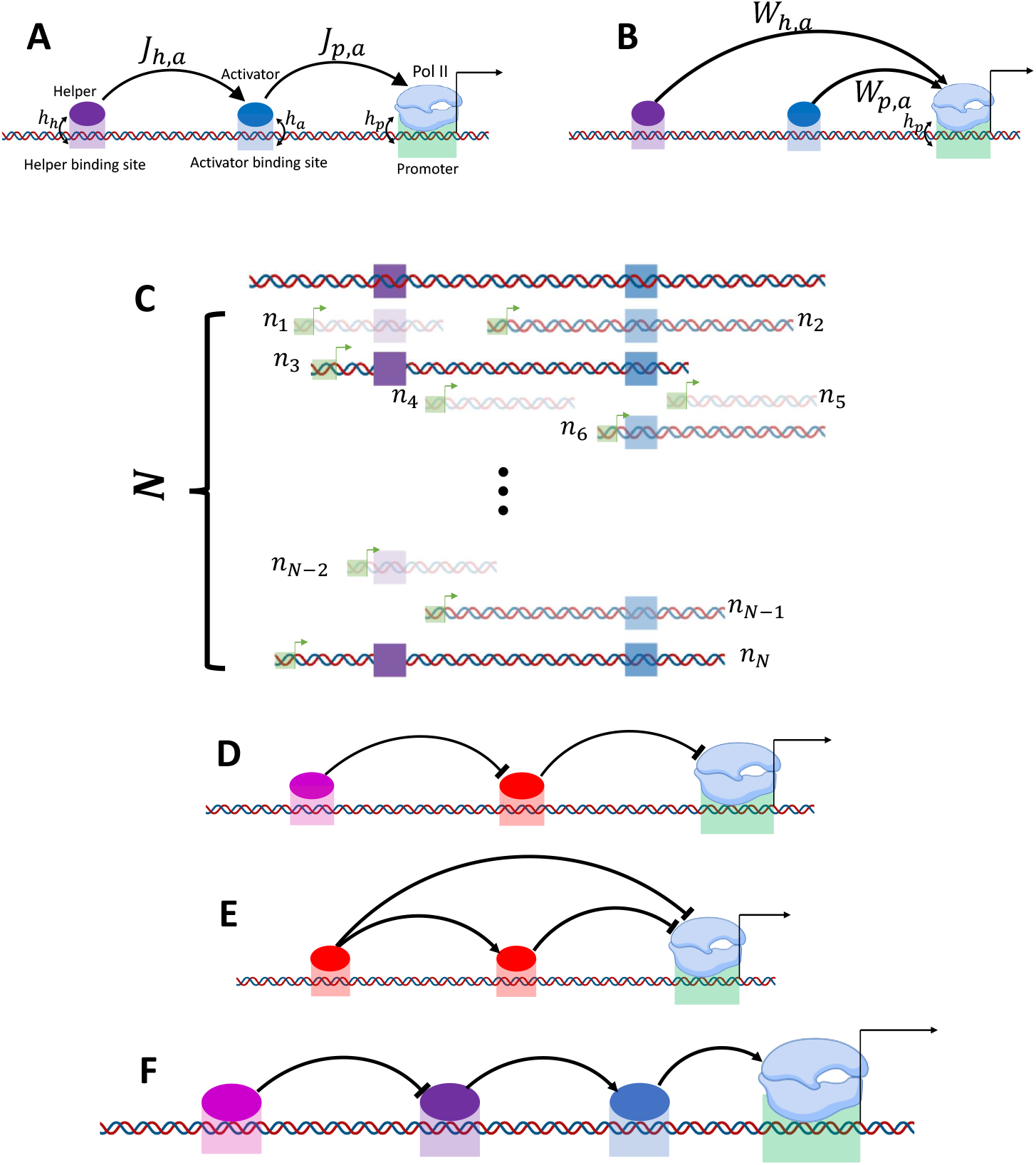
A) Schematic of a helper-activator enhancer. Shown are the non-zero interaction energies between the TFs and Pol-II for an enhancer bound by an activator (blue) and a helper (purple). B) A diagram showing the effective interaction energies to be inferred by fitting a mean-field model of transcription to the data. C) Illustration of the data collected in a typical STARR-seq experiment for the helper-activator enhancer shown in (A). The data consists of *N* sequence tiles that overlap the region, each with a measured number of transcribed mRNA, *n*_*i*_. Tiles containing both helper and activator sites exhibit higher activity than those with only the activator site, which in turn are more active than tiles with a helper site alone. Tiles lacking any binding site show the least activity and are represented with greater transparency. In D-F, we show other synthetic enhancers for which we simulated data and applied our inference method on. D) A system involving an inhibitor (pink) and a repressor (red). E) A system comprising two repressors, where the left repressor acts as a supportive element for the right repressor. F) A system incorporating an activator, a helper, and an inhibitor of the helper.

Given a set of interaction energies between TFs and Pol-II, as well as the binding site energies for various factors, and assuming the system is in thermodynamic equilibrium we can calculate the probability of Pol-II being bound as

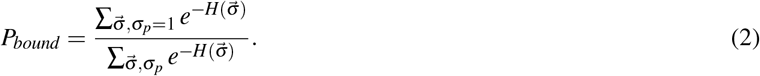

This model is equivalent to the one described in [16], which employed a combinatorial approach to estimate *P*_*bound*_.

Given the equilibrium probability of Pol-II being bound, for simplicity, we assume that the production of mRNA follows a Poisson process with a rate being proportional to *P*_*bound*_. The expected number of transcribed mRNA for this process is *µ* = *P*_*bound*_*r*_0_, where *r*_0_ reflects the maximal expected mRNA output as measured in the experiment for a single enhancer.

### 3.2 Generating synthetic STARR-seq data

Here we describe our method for generating synthetic STARR-seq data on which to test the inference algorithm. To simulate a STARR-seq experiment for an enhancer region of interest, we first start by generating a random set of *N* overlapping sequences known as tiles, that cover the region (see Fig. 1(C)). The positions of the tiles are drawn uniformly from the region of interest, and their lengths are drawn from a normal distribution with the mean length *µ*_*L*_, and standard deviation *σ*_*L*_. The enhancer region is set to have a defined number of TF binding sites with fixed positions and widths, *x*_*i*_ and *l*_*i*_ respectively, each with their own site-energy *h*_*i*_ and interactions with other sites, given by *J*_*i, j*_. As seen in Fig. 1(C), a given tile may overlap all, some or none of the TF binding sites. If a tile does not completely overlap the entire binding site, that site is not considered to be in that tile. This is how the width of the binding site enters the model, by determining whether a site is in a tile or not. In the STARR-seq assay, all tiles possess the same Pol-II binding site, which we model with corresponding site energy, *h*_*p*_, and interactions with the TF sites, *J*_*p, j*_.

For the *i*-th tile, the probability of Pol-II binding, 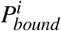, is calculated using Eq. 2 for the binding sites contained within that tile, whose expected number of mRNAs is 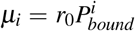, given some value for the maximal expression level, *r*_0_. We then generate a random sample of mRNA counts (one for each tile), {*n*_*i*_ }, from a Poisson distribution given the corresponding *µ*_*i*_. The amount and quality of the synthetic data is controlled by the number of tiles, *N*, and *r*_0_, which is related to both the experiment duration and sequencing depth.

### 3.3 Inferring the model parameters from data

Our goal is to infer the number, locations, and widths of potential TF binding sites within an enhancer, along with their interaction energies with Pol-II, and the Pol-II site energy, *h*_*p*_, given only a set of observed mRNA counts from a set of sequence tiles that overlap the region of interest (see Fig. 1B). Given that the only information we have is the mRNA counts {*n*_*i*_}, it is not possible to infer the full interaction matrix, *J*_*i, j*_. This is because the system is underdetermined, since we do not have any measurements of the correlations between TF binding sites, and the experiment is only capturing correlations between the TFs and Pol-II. To overcome this challenge, we instead try to infer the parameters of a mean-field model for the Hamiltonian given by Eq. 1. Using the mean-field approximation, the probability of a Pol-II being bound to the *i*-th tile, 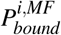, is given by

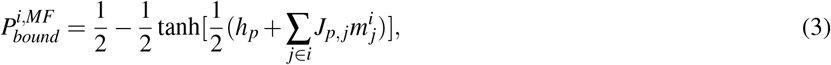

where 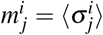 is the average occupancy of binding site *j* on tile *i*, and the sum over *j* is just over those binding sites that are wholly within tile *i*. For the inference model we are considering, we will assume that the 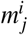 is not measured or known. Furthermore, we will make the simplifying assumption that 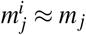, which is the average occupancy of the site over all tiles. By doing so, we will define an effective interaction weight between Pol-II and the *j*-th TF binding site as, 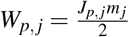. Thus Eq. 3 becomes

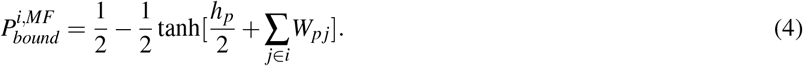

We assume that the observed mRNAs are drawn from a Poisson distribution, 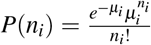, where the expected number of mRNA from a tile is 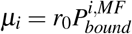.

For a model with a fixed number of TF binding sites, the parameters, *θ*, to be inferred are their locations, *x*_*i*_, widths, *l*_*i*_, effective interaction energies with Pol-II, *W*_*p, j*_, and the Pol-II site energy, *h*_*p*_. The parameter *r*_0_, which sets the maximum expected amount of mRNA output, is not inferred. We set it to be either the *r*_0_ that was used when making the synthetic data, or to be a value close to the highest mRNA counts (i.e. assumes the most highly expressing tile has *P*_*bound*_ ≈ 1). Given a set of values for all of these parameters, the log-likelihood of the data (i.e. the observed mRNA counts, {*n*_*i*_}) is given by

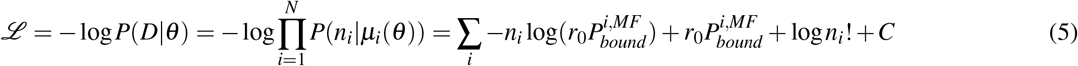

where we have taken *P*(*n*_*i*_|*µ*_*i*_(*θ*)) to be the Poisson distribution, and 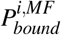 is the probability of Pol-II being bound corresponding to the *i*-th tile, which is computed using Eq. 4.

To determine the optimal parameters that minimize the log-likelihood function, we use Monte Carlo optimization. This algorithm involves iteratively updating the values of the parameters, where during each iteration a randomly selected parameter is modified. This modification entails adding *a*_*k*_*η* to the parameter, where *a*_*k*_ represents the step amplitude of the *k*-th parameter, and *η* is a random number drawn from a Gaussian distribution with a mean of zero and a variance of one. If the new value of the selected parameter reduces the log-likelihood function, it is accepted. Otherwise, it is accepted with a probability of 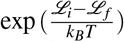, assuming *k*_*B*_*T* = 1. It is worth mentioning that although the width and position of a binding site are discrete variables measured in base pairs, for computational simplicity we treated these as continuous parameters, similar to our energy parameters. We update *a*_*k*_ every specific number of iterations (*N*_*I*_). Following every *N*_*I*_ iterations, the algorithm computes the success rate (*S*_*R*_), which represents the number of accepted moves over *N*_*I*_. In order to maintain the success rate around 0.5, 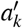 is updated as 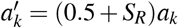. This helps prevent getting trapped in local minima, or making drastic jumps in the energy landscape. If the success rate exceeds 0.5, *a*_*k*_ increases, leading to a decrease in 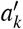, thereby reducing the success rate, and vice versa. The algorithm is executed for several thousand iterations, allowing the likelihood function to converge towards a stable value. Upon completion, the algorithm identifies the parameter set θ^*^ corresponding to a minimum of the log-likelihood function. It is important to note that we adjust the number of iterations proportionally for systems with more binding sites, which have more parameters. During the search for the best parameters, all parameters are freely adjustable except for the binding site width, which is limited to a range of 5 to 15 base pairs. This limitation guarantees that the identified binding site widths remain within a reasonable range.

The value of the optimal parameters can depend on their initial values. So for a given dataset, the code is run 15 times with different initial values, and the parameters that give the lowest log-likelihood are chosen. To initialize the energies, we sample initial values from a Gaussian distribution with a mean of zero and a variance of one. For the initial position of sites, we randomly select a number between the location of the starting point of the region plus 15 base pairs and its ending point minus 15 base pairs. Additionally, for the width of the sites, we choose random values within the range of 5 to 15 base pairs.

### 3.4 Selecting between models with different number of binding sites

Since we do not know how many binding sites are present in an enhancer, the task involves comparing models of different complexity. We use the log-likelihood function (*ℒ*), to compare models. Models with more binding sites typically show lower log-likelihood than those with fewer, however, at some point the gains diminish with increasing complexity. To select a model that is complex enough to capture the necessary details without overfitting, we find the point in the plot of *ℒ* versus number of binding sites where *ℒ* ceases to decrease significantly, and choose the corresponding model—this point is known as the ’elbow’ or ’shoulder’ of the curve.

To determine the elbow of the curve, we first normalize the data. Then, we calculate the distance of each data point from the line *y* = *x*. Subsequently, we select the model that corresponds to the greatest distance from *y* = *x* [21]. In Section 4.2, due to the noisy behavior of the *ℒ* curve for each region, we fit a suitable polynomial to each curve.

### 3.5 Bin definition

When assessing a binding site’s position within a region, slight variations in its precise placement may not impact the log-likelihood function, as it may not change the group of tiles covering the site. Consequently, while our model can confirm the presence of the binding site within that specific area, its exact position remains indeterminate. We refer to these areas as “bins,” which are defined by initially marking the beginning and ending of every tile across the region. Each bin is then defined as the interval between two consecutive markers, resulting in variable bin length and a unique bin set for each dataset. Therefore, in the fitting process, we assign the energy weight of a found site to all the base pairs of the bin where the site is found.

Since we generated ten different random configurations of tiles for each enhancer architecture in Section 4.1, each base pair is assigned ten energy values. We report the average of these values as energy weight. For the experimental datasets in Section 4.2, where there is only one dataset per region, we do not take an average.

### 3.6 Quantifying classification of known binding sites

To evaluate our model’s ability to infer a known set of sites, we treat it as a binary classifier. As such, we assign 1s to base pairs with an inferred binding site and 0s to all other base pairs. Furthermore, we construct a ground truth binary vector for the actual system, allocating 0s and 1s to the base pairs where a known binding site is absent or present, respectively. By comparing the inferred vector to the ground truth, we can categorize each base pair as true-positive, false-positive, true-negative, or false-negative. This enables us to calculate various classification metrics such as accuracy, precision, recall, etc.

To quantify an inferred model’s classification ability, we calculate the *f*_1_-score, which is defined as:

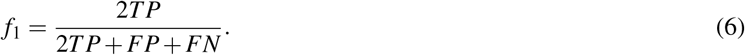

It ranges between 0 and 1. A higher *f*_1_ value indicates fewer false negatives and false positives simultaneously, thereby reflecting the reliability of the prediction.

### 3.7 Including degeneracy of tiles

In a STARR-seq experiment, a given tile, *i*, may occur multiple times within the library. This degeneracy can be determined from sequencing, providing a count *L*_*i*_ for the number of occurrences within the library. Each copy of this tile produces an associated mRNA count, which is assumed to be generated from a Poisson process with mean, 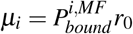, as explained in previous sections. However, the experimental measurement corresponds to the sum of counts across all copies. For example, for the *i*-th tile, the total count 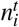 is calculated as 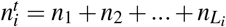, where *n*_*j*_ represents the count of the *j*-th copy. Since *n*_*j*_ follows a Poisson distribution with a mean of *µ*_*i*_, 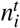 also follows a Poisson distribution with a mean of *M*_*i*_ = *L*_*i*_*µ*_*i*_. This makes a slight modification to the equation for the log-likelihood function, by replacing *µ*_*i*_ with *M*_*i*_ in Eq. 5.

### 3.8 Motif identification

Given the DNA sequence of an enhancer region, we can identify possible locations of binding sites by using known motifs for a given set of TFs. For the *k*-th TF, we have a position probability matrix, 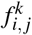, which gives the probability of observing *i*=A,C,T,G at position *j* in the motif. We scan each motif along the forward and reverse strands of a sequence and calculate a binding score at each location and direction using

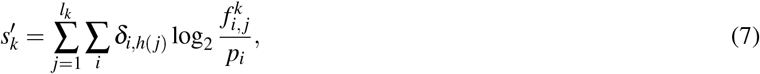

where *l*_*k*_ represents the length of the motif, *δ* is the kronecker delta, *h*(*j*) shows the nucleotide type at position *j* (which can be A, T, C, or G), and *p*_*i*_ is the background probability for each nucleotide, set at 0.25 for all types. The final score of the motif is determined using the following equation:

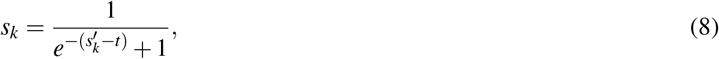

where *t* is a threshold, which is set to 5 for all TF motifs. A higher value of *s*_*k*_ indicates a greater likelihood of binding of the *k*-th TF at that specific location in the sequence.

TF motifs MA0007.2.AR, MA0148.3.FOXA1, and MA0901.1.HOXB13 [22], were chosen for the AR enhancers regions as they are known to be involved in AR enhancer regulation [23–25].

### 3.9 Self-transcribing active regulatory region sequencing (STARR-seq)

STARR-seq was done as previously described [26]. Briefly high-confidence androgen receptor (AR) binding sites (n=4139) were tested in an adenocarcinoma prostate cancer cell line (LNCaP). To identify androgen-driven enhancer activity cells were grown in charcoal stripped serum for 72 hours and then treated with EtOH of 10 nM dihydrotestosterone for 6 hours. Libraries were prepared as described and processed using a standardized pipeline.

### 3.10 Chromatin immunoprecipitation followed by sequencing (ChIP-seq)

Chromatin immunoprecipitation followed by sequencing (ChIP-seq) Publicly available ChIP-seq data (GEO GSE83860, GSE83860, GSE94682) was used to identify AR, FOXA1 and HOXB13 binding sites. Data was processed using a standardized pipeline [27].

## 4 Results

### 4.1 Inferring binding sites in synthetic enhancers from simulated STARR-seq data

We now detail the results of applying our inference method (see Methods) for identifying the transcriptional regulation of several simple enhancer architectures [17]. The first architecture we will consider is shown in Fig. 1(A), which consists of two binding sites for an activator and helper transcription factor, respectively. The role of the activator in this system is to increase the binding probability of Pol-II, and the helper, further stabilize the binding of the activator. Our inference method is learning a mean-field representation, and as shown in Fig. 1(B), the helper now has an effective interaction with Pol-II that captures some of the correlations it has with Pol-II binding through its interaction with the activator. Given locations and widths for the binding sites of the helper and activator along with their interaction energies (see Supplementary text for actual values), we then proceed to simulate data from a STARR-seq assay (see Methods). As shown in Fig. 1(C), the enhancer region is partitioned into a set of *N* overlapping sequences, known as tiles. It is assumed that the same promoter binding site is present in all tiles. A given tile may overlap both activator or helper sites or one or none at all. For the *i*-th tile, we use the fully interacting model (Fig. 1(A)) to calculate the equilibrium probability of Pol-II being bound, 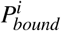, from which we then generate a random number of mRNA from a Poisson distribution with mean, 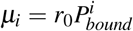, where *r*_0_ sets the maximum number of expected mRNA counts in the STARR-seq assay. The simulated data yields a set of mRNA counts for each tile, *n*_*i*_, as shown in Fig. 1(C).

Fig. 2(A) represents the results from fitting our mean-field model to ten distinct synthetic datasets for a 700-base-pair enhancer with a helper site and an activator site (Fig. 1(B)), where the number of tiles, *N*, and maximal expression, *r*_0_, were fixed, but where the locations of the tiles differed. The top subplot of the figure shows the position and width of the helper and activator sites (see Supplementary Material for precise values) over the ∼ 1500-base-pair regions spanned by the overlapping tiles. The blue curve represents the average number of mRNAs at a given base pair across all tiles, indicating the transcriptional activity within the enhancer region.

**Figure 2.**
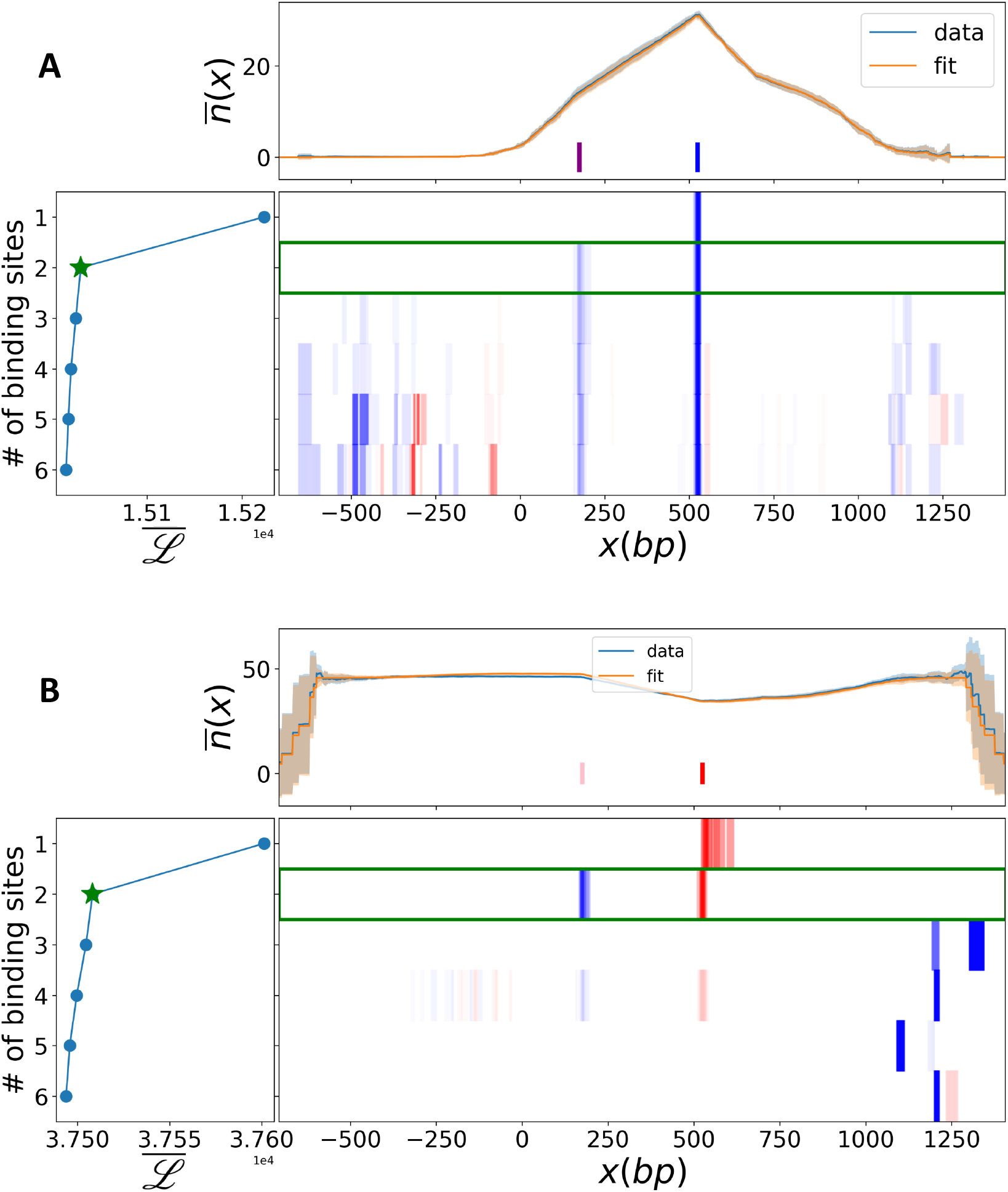
Figures A) and B) represent the helper-activator enhancer and the repressor-inhibitor enhancer, respectively. Colored bars in the upper subplots show the known position and width of binding sites, where purple and blue in the helper-activator system show the helper and activator sites, and pink and red in the repressor-inhibitor enhancer show the inhibitor and repressor sites, respectively. The average number of produced mRNA over all tiles spanning a given base pair is depicted in light blue for the data and orange for the fitted mean-field model with two binding sites, with the shade indicating the standard deviation from the mean across ten different tile configurations. The lower-left subplots depict the average log-likelihood, *ℒ*, against the number of binding sites across ten configurations of tiles, with the green star marking the shoulder of the curve. The color coding in the lower-right subplots represents the interaction energies as inferred by the mean-field model. Each row shows the average weight of inferred sites at each base pair for the models with a given number of sites to infer. In these, red and blue represent positive and negative weights, respectively, indicative of a base pair’s repressive or activating role. The intensity of the color corresponds to the absolute magnitude of the energy. This subplot was created by assigning the energy value of a found binding site to every base pair within the bin where this binding site is located. The green rectangle shows the optimal model based on the shoulder of the *ℒ* curve. Fixed values of *r*_0_ = 50 and *N* = 400 were used to simulate both systems.

The leftmost plot of Fig. 2(A) shows the average log-likelihood over the ten different datasets as a function of the number of sites in the model. We identified the shoulder of *ℒ* for each dataset separately, and in all cases, the model with two sites was located at the shoulder. The orange curve shows the average number of mRNAs at a given base pair across all tiles, estimated by the expected count of each tile 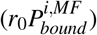 from the selected model (two-site model). The corresponding site locations for model with two sites, align well with the known locations of the helper and activator. Since we are fitting a mean-field model of transcription, each site has an effective interaction with Pol-II. As can be seen, the activator site has an effective attractive interaction with Pol-II (dark blue), but so does the helper. In the actual helper-activator architecture, there was no explicit interaction between the helper site and Pol-II (see Fig. 1(A)), but because the helper increases the output when in tandem with the activator, it effectively acts like a weaker activator (lighter blue). Thus, the inferred model is able to assign effective transcriptional activity to the helper and activator sites in addition to finding their correct locations.

Additionally, we conducted similar analysis for the three other enhancers depicted in Fig. 1 (D-F). In the repressor-inhibitor system (Fig. 1(D)), the repressor (red site) inhibits Pol-II from binding to its promoter, and the inhibitor (purple site) weakens the binding of the repressor to its own binding site. Although the inhibitor does not have any direct interaction with Pol-II, it effectively assists the binding of Pol-II by blocking the repressor. As for the first enhancer, we simulated STARR-seq data for the repressor-inhibitor enhancer and the results are presented in Fig. 2(B). The top plot shows the transcriptional output as a function of position within the region spanned by the tiles. As can be seen the output is overall roughly constant over the region, except for tiles that possess only the repressor site, where there is a drop in output. We inferred models with different number of binding sites, and found that the shoulder in log-likelihood was at two sites (left plot). The average weights and positions for two-site models are shown in the bottom map. The effective mean-field model with two sites finds that the repressor has a repulsive interaction with Pol-II (red), but that the inhibitor of the repressor has an effective attractive energy with Pol-II (blue). This was as expected, and shows that the mean-field model can provide functionally correct assignments to bound factors.

The results for the other two enhancer architectures (Fig. 1 (E-F)) are shown in Fig. S1. The inference method correctly found two sites for the dual repressor architecture (Fig. 1(E)) and that they both functionally act as repressors of Pol-II binding. For the last simple architecture (Fig. 1(F)) that consists of three sites, an inhibitor, helper and activator, the method identified the three-site model as the shoulder of the log-likelihood, where the helper’s inhibitor is effectively functioning like a repressor of Pol-II, and the helper is, as before, functioning like an activator. Thus for these simple architectures, the chosen models had the same number of sites as in the original enhancers and made functionally correct assignments to the sites.

#### 4.1.1 Impact of data quality on the inference process

We now explore how data quality impacts the inference method. Data quality is controlled by two parameters: *N*, the number of tiles covering the enhancer of interest, and *r*_0_, the expected maximum measured mRNA output from a tile, that depends on sequencing depth as well as duration of the transcriptional protocol.

For the four previously discussed enhancers, we used our inference method on ten datasets per enhancer created using different *r*_0_ and *N*, and in each case, chose the model at the shoulder of the log-likelihood. In Fig. 3(A), we show how the inference of the helper-activator system is improved by increasing the number of measured tiles, *N*. Specifically, it illustrates how the uncertainty in locating the binding sites improves with *N*, as a greater number of tiles provides greater resolution.

**Figure 3.**
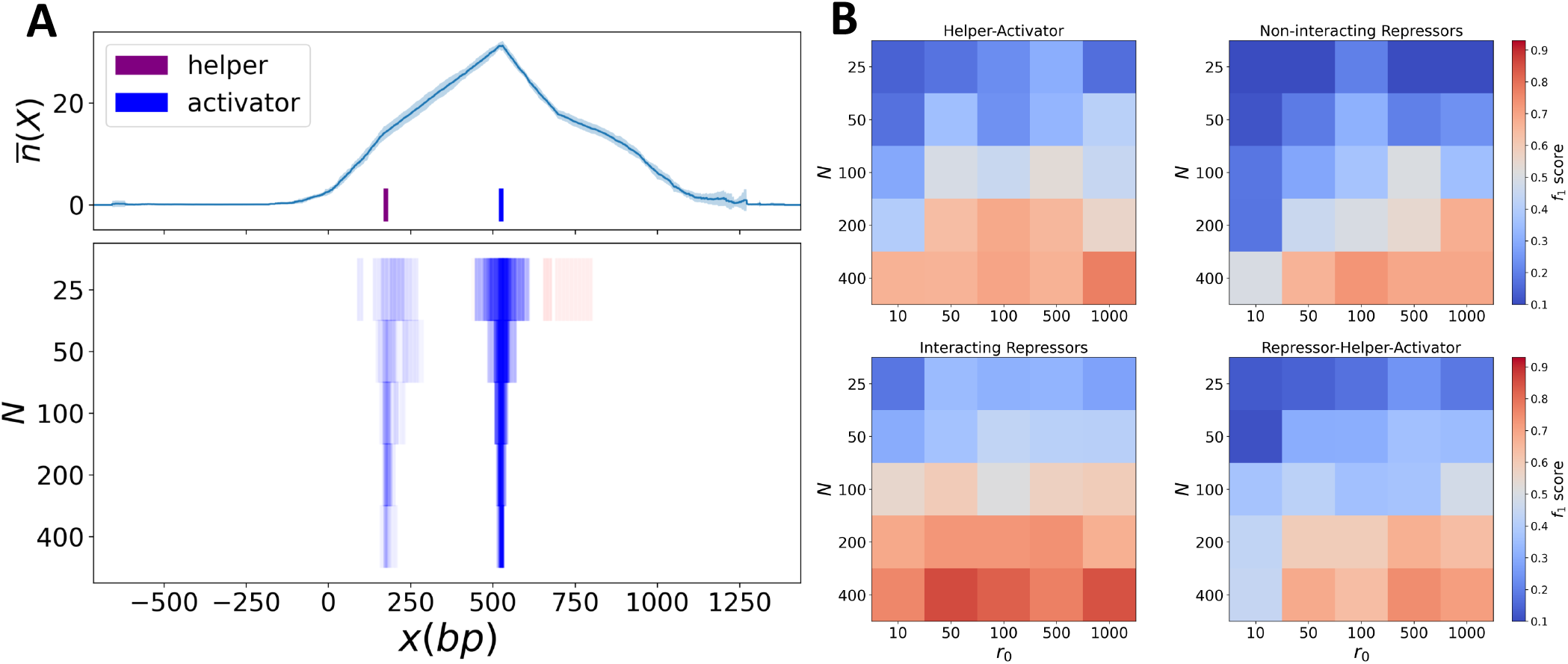
Impact of data quality on the inference method. Fig. A, the upper subplot, represents the average mRNA output over tiles of the helper-activator system for the case where *N* = 400 and *r*_0_ = 50. The lower subplot shows inferred binding sites for the select models at different number of tiles, *N*, and with *r*_0_ = 50. Each row shows the average weight over ten different configurations of *N* tiles. Blue and red represents negative (activating) and positive (repressing) role, respectively. As evident, the efficacy of inference increases with *N*. B) Heat maps showing the quality of classifying the original enhancer (*f*_1_-score) as a function of the number of tiles, *N*, and expected output level *r*_0_, for the four enhancer architectures shown in Fig. 1(A,D-F).

As described in Methods, we calculate the *f*_1_-score between the locations of the inferred sites in the model and those of the original enhancer. When the number of tiles is low, the best model tends to over predict sites (see Fig. 3(A), for *N* = 25, 50), leading to low *f*_1_-score. However, as *N* increases, the false-positives decrease and the *f*_1_-score improves. Fig. 3.(B) shows color coded average *f*_1_-score for enhancers shown in Fig. 1. As can be seen, in all cases *f*_1_-score increases with *N*. Similar to the helper-activator system, the other enhancers also showed an improvement in *f*_1_-score as the number of tiles were increased. The effects of amount of measured mRNA, through *r*_0_, shows more of a transition where the *f*_1_-score exhibits a relatively rapid improvement as *r*_0_ increases, particularly beyond a value of *r*_0_ = 50. Thus for these simple enhancer architectures, our findings suggest that having *N >* 100 and *r*_0_ *>* 50 can lead to consistently good inferences of the true sites within enhancers.

#### 4.1.2 Enhancers with more complex regulatory architecture

We previously examined enhancers with a small number (≤ 3) of binding sites and here we consider enhancers made up of a larger number of sites with more complex regulatory architectures.

As we wish to design a complex enhancer, we chose to randomly assign the site energies and the energetic interactions between TFs and between TFs and Pol-II for our chosen number of TF sites. Given the random interactions, could the inference method infer an effective model that describes the output of such a complex enhancer?

The output of the enhancer (the blue curve in Fig. 4(A) upper subplot) shows several levels of expression, with the largest occurring at the right side of the sequence. We can gain some insight into the function of each site by looking at the interaction energies of each binding site with Pol-II shown as the orange squares in Fig. 4(B). Here the 3^*rd*^ and 7^*th*^ sites have attractive (activating) interactions with Pol-II, while all other sites have positive (repressive) interactions. Even further insight can be found by looking at the effective interactions, 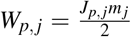 where *m*_*j*_ is the expected occupancy of the *j*-th site which we calculate from the full Hamiltonian (Eq. 2) (see blue dots in Fig. 4B). Now we see that binding sites 1, 4, and 6 have *W*_*p, j*_ ≈ 0, highlighting that they are likely not bound, and hence should not have any functional role. With these observations in mind, we now look at what our inference method determined for the mean-field model of this enhancer.

**Figure 4.**
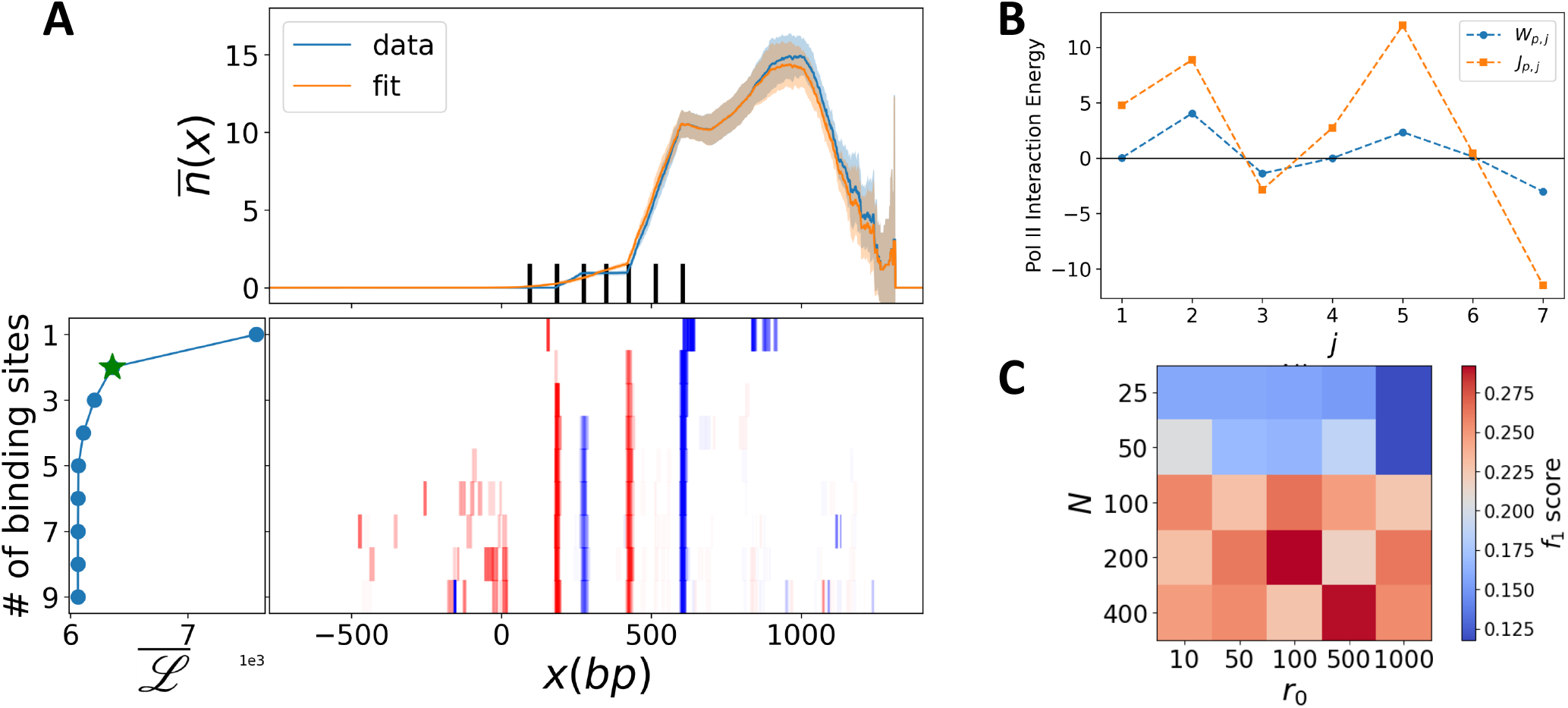
A) Black bars in the upper subplot show the position and width of binding sites (please see Supplementary Material for the exact values of locations and widths). The blue curve represents the average mRNA count over tiles of the system for the case where *N* = 400 and *r*_0_ = 50. The orange curve shows the same quantity estimated by the expected count of each tile 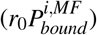 from the select model, which here is the two-site model. In these, red and blue represent positive and negative weights, respectively, indicative of a base pair’s repressive or activating role. B) The figure shows the interaction energy of each binding site with Pol-II. *J*_*p, j*_ is the interaction energy (orange dots), while 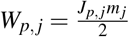 is the effective interaction energy with the *m*_*j*_ calculated from the full model (Eq. 2), but not the mean-field model. C) A heat map showing the quality of classifying the original enhancer (*f*_1_-score) as a function of the number of tiles, *N*, and output level *r*_0_.

The lower subplot of Fig. 4(A) illustrates found sites for models with different number of binding sites. As can be seen, the mean-field model at the shoulder of the *ℒ* curve (leftmost subplot of Fig. 4(A) has found three sites (sites 2, 5, and 7) out of the seven original binding sites (in each dataset, one of these three sites is missing). This makes sense in light of our description of the binding sites above, where sites 1, 4, and 6 are not bound (Fig. 4(B)); having no function, and hence not inferable, and site 3 has a weaker binding energy. The effective interactions of the three functional sites are consistent with their known interactions with Pol-II as either activating (site 7) or repressing (sites 2 and 5). Thus the selected model has inferred only the stronger functional sites. This is an important result, as models which are based on motifs, either known or de novo, could have sites at the non-functional locations, potentially leading to a poor explanation of the output of such an enhancer.

We also looked at how data quality influenced the ability of the inference scheme to infer the known binding sites. The *f*_1_-score is plotted in Fig. 4(C) as a function of the number tiles, *N*, and the expected maximal mRNA output, *r*_0_ (also see Fig. S2(C)). As expected, it increases with the number of tiles, however the effect of varying *r*_0_ does not show any obvious trend. Moreover, the maximum score is around 0.4, which is lower that the maximum score for previous simpler enhancers. This is the result of using all seven known sites as the ground truth, some of which are not functional (which may not be known a priori, if just annotating motifs) and hence not inferable, leading to a fixed number of false negatives.

Furthermore, we considered another enhancer with seven TF binding sites (see Fig. S2). As illustrated in Fig. S2(A), the average expression level on the left side of this enhancer is higher compared to the right side, which is nearly inactive. This observation aligns with the energetic interactions of sites with Pol-II, *J*_*p, j*_ (Fig. S2(B)), where site 4 to 7 exhibit positive interaction energy, indicative of repressing role. Site 1 also has positive *J*_*p*,1_, however, it is located next to two activators with negative *J*_*p, j*_, which can weaken its repressing role. Fig. S2(A), the lower-right subplot, shows the inferred binding sites for models with different number of sites. As can be seen, only sites 3 and 4 out of the seven sites have been inferred by the mean-field model, which is consistent with the number of sites at the shoulder of *ℒ* curve in the lower-left subplot. Again, this can be explained by looking at Pol-II’s interaction energies, Fig. S2(B), which shows the known values of *W*_*p, j*_ calculated using the full model (Eq. 2): two of the non-identified sites are site 5 and 7, whose *W*_*p, j*_ is around zero, indicating their inactivity and negligible impact on transcription. Site 6 has a positive *W*_*p, j*_, suggesting a repressive role, but the average number of tiles’ mRNA per base pair (the blue curve in Fig. S2(A)) at this site is around zero or equivalently, *P*_*bound*_ ≈ 0, which means the site is inactive and cannot be identified. *W*_*p, j*_ is greater than 0 for site number 1, however, it cannot be inferred because of a different reason: it is located next to site number 2, with similar weight in absolute terms, i.e. |*W*_*p*,1_| ≈ |*W*_*p*,2_ |. There are two kinds of tiles covering site 1; those including only site 1, where *P*_*bound*_ = 0 due to *h*_*p*_ = 10 (refer to Eq. 4), and those covering both sites 1 and 2. For the latter, the effect of site 1 on *P*_*bound*_ is nearly offset by site 2. Even though *W*_*p*,2_ *<* 0, site 2 isn’t inferred either. The bump around *x* =−150*bp* is attributed to site 3 because all tiles covering that area, along with site 2, also cover site 1, and we know that sites 1 and 2 cancel each other’s effect. Only tiles that cover site 2, or sites 2 and 3, or sites 2, 3, and 4, or sites 2, 3, 4, and 5, can aid in identifying site 2. However, all tiles covering sites 1 and 2 simultaneously have zero output, suggesting there are no sites at the location of site 2. Therefore, the model has merged sites 2 and 3, both activators, at the location of site 3.

### 4.2 Inferring regulatory sites from experimental STARR-seq data

Here we use our inference method on experimental STARR-seq data [20] that measured the transcriptional activity of androgen receptor (AR) bound regions in the human genome. AR is a transcription factor, that when bound by androgens, enters the nucleus and regulates the transcription of its target genes. Its aberrant activity is a key driver of the development and progression of prostate cancer [28]. In the published experiment, 4139 AR bound regions were tested for activity, and 286 were found to be differentially active when induced by hormone, Dihydrotestosterone (DHT). We applied our inference method on 36 regions of the AR bound regions, which are AR-driven enhancers, to compare our inferred regulatory sites (we now use the term regulatory site, as it may be possible for a sequence to regulate transcription yet not be directly bound by a TF) with the ChIP-seq available data and binding score of three TF motifs (AR, FOXA1, and HOXB13) known to be involved in AR enhancer regulation.

We show the results of our inference method on one of the regions in Fig. 5(A-D). Fig. 5(B) shows the result of our fit to the mRNA output of 34 wild-type sequence tiles that overlap the full 700 base pair region in the presence of DHT using models with a range of regulatory sites from 2 sites to 29 sites. As in our prior analysis on synthetic enhancer architectures, blue sites correspond to activating sequences while red ones correspond to repressive function. The model made up of 10 regulatory sites corresponds to the shoulder of the log-likelihood. However, across all models, it can be seen in Fig. 5(B) that there is a general consensus of the strong activating sites near 410 bp, peaks in the ChIP-seq score plot (Fig. 5(C)), and the location of the AR motif in the binding score plot (Fig. 5(D)). Other activating sites (50, 90, 180, 250, 310, 610 bp) were also found in this AR-bound region, suggesting other important functional elements. For instance, based on the infered model, the site located at x=610 bp is a strongly activating site (see Fig. 5(A,B)), where none of the three motifs with a high binding score is present there (see Fig. 5(D)). This could be an example of an important functional site that can be missed in global fitting approaches.

**Figure 5.**
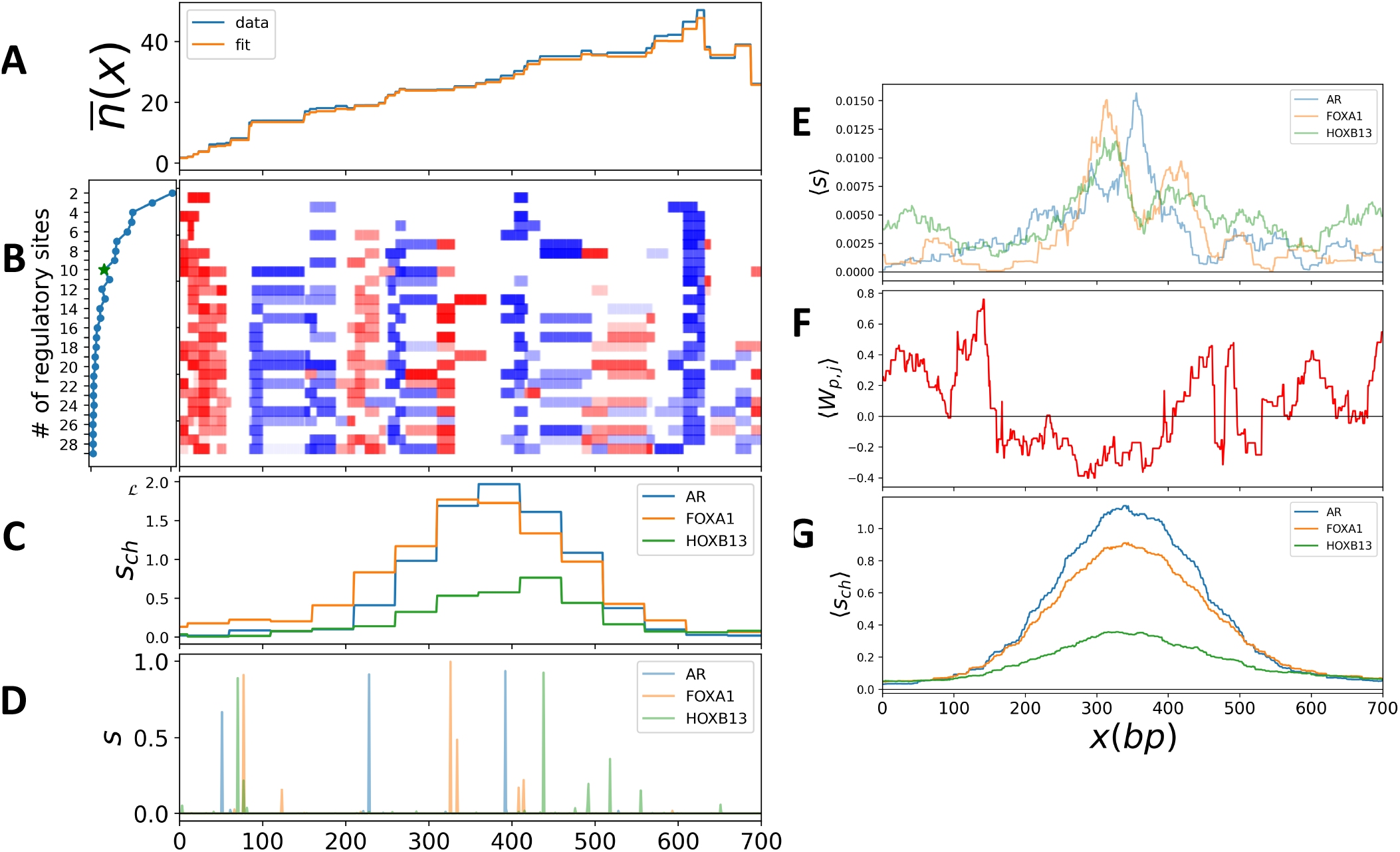
Fig. (A) represents the average mRNA output over tiles for both the data in blue and the fitted model with 10 regulatory sites (shoulder of *ℒ* curve). B) Inferred regulatory weights for models with differing numbers of regulatory sites. In these, red and blue represent positive and negative weights, respectively, indicative of a base pair’s repressive or activating role. Fig (C) and (D) show ChIP-seq score and binding scores, respectively, for known motifs in the sequence of the region ’peakno-2415-ARBS’ and its reverse complement. E) Average binding score of the known motifs over all 36 AR-bound regions that we applied our inference method to. F) Average of inferred weights of the selected model in each region (shoulder of log-likelihood, see Fig. S3). G) Average ChIP-seq score for the three known motifs across all 36 regions.

We summarized the results for all the 36 regions in the next three plots (Fig. 5(E-G)). Fig. 5(E) illustrates the average binding score for the three known motifs (AR, FOXA1, HOXB13). As can be seen, there is a peak around the middle of the plot, indicating the higher concentration of the motifs around the center of regions. Fig. 5(G) presents the average ChIP-seq score, where the peaks confirm the higher binding activity of these TFs around the middle of the regions. Fig. 5(F) shows the average inferred weights for these 36 regions at the shoulder of the log-likelihood curve (as shown in Figure S3). A comparison between these three plots reveals a good general agreement in their trends, the location of the peak in the top and lower plots aligns with the location of the negative activating regions in the middle. Despite the relatively low number of tiles per region (on average 29 tiles per region), there is general agreement about the strength and location of transcriptional regulatory elements that is encouraging.

## 5 Discussion

In this paper, we have introduced a method for identifying putative regulatory sites within an enhancer sequence using data from a STARR-seq assay that measures the mRNA output of overlapping sequence tiles that cover the enhancer. We modeled the transcriptional output of each tile by using a mean-field model to calculate the probability of Pol-II being bound given a fixed set of regulatory (TF binding) sites. We modeled the binding of Pol-II using an Ising-type model, enabling us to simulate the transcriptional output of the STARR-seq assay. This approach allowed us to determine the locations, widths, and interaction energies of binding sites with Pol-II, representing their effective role in transcription within a region by minimizing the log-likelihood of the data given our model.

We first tested our model on simulated STARR-seq data for some simple enhancer architectures where we specified the locations and functions of each binding site. In all cases, the inferred model was able to find the location of the known sites, and assign them an effective interaction with Pol-II that was consistent with their known function (e.g. a repressor of a repressor effectively functions like an activator, as was inferred by the model). We then explored the impact of data quality through the number of tiles, *N*, and the expected number of mRNAs, *r*_0_, on our inference results. We found that having more tiles spanning a region led to greater resolution of the inferred sites and greater classification of the known sites, while the expected number of mRNAs has less pronounced effects. Additionally, we applied our model to more complex enhancer architectures and demonstrated its effectiveness at identifying regulatory sites that had a functional role in transcription. In particular, there were known sites that were not bound, and the model correctly inferred that there were no regulatory sites at those locations. This is one of the strengths of this approach over methods based on known or discovered motifs that would then try and fit the assay output using identified motif sites that could be non-functional. These complex architectures also highlighted how dissecting nearby repressor sites can be challenging, since there is little variation in the output on which to try and assign function.

We then applied our method to experimental STARR-seq data for AR-driven enhancers measured in a prostate cancer cell line. Although the data for each region was insufficient for a robust fit, we observed a promising agreement between the effective function of the inferred sites and an independent experimental dataset (ChIP-seq data). Moreover, the locations of known motifs overlapped locations of identified sites by the model. To reduce errors and enhance the accuracy of our method, we recommend increasing the number of tiles per region in future studies.

Thus our model shows promise at being able to identify important regulatory elements within an individual enhancer. Here we have only used it to fit the data for a single region at a time, therefore allowing it to find potentially local and novel regulatory elements that may be missed in a global fit. One could imagine extending it to a global fit by specifying the regulatory sites to be associated with a specific factor, thereby learning its average functional effect genome wide. Such a global fit could potentially discover the optimal set of regulatory factors that capture the observed transcriptional data.

Other extensions involve generalizing some of the assumptions that we made. We assumed that the production of mRNAs follows a Poisson process, a choice that simplifies the model. A more realistic approach for future improvement could be the incorporation of a negative binomial process, which better represents transcriptional variability. Additionally, we initially assumed that the average occupancy of a site is constant regardless of the different tiles covering it. However, this may not accurately reflect biological reality, as interactions between sites can influence their occupancy. An enhancement to our model could involve relating the occupancy of a site to the presence of other sites within the same tile. Nevertheless, we think that this method, as it currently stands, can serve as an additional tool that researchers can use to decipher the transcriptional logic for their system of interest.

## 6 Supporting Information

**S1:**
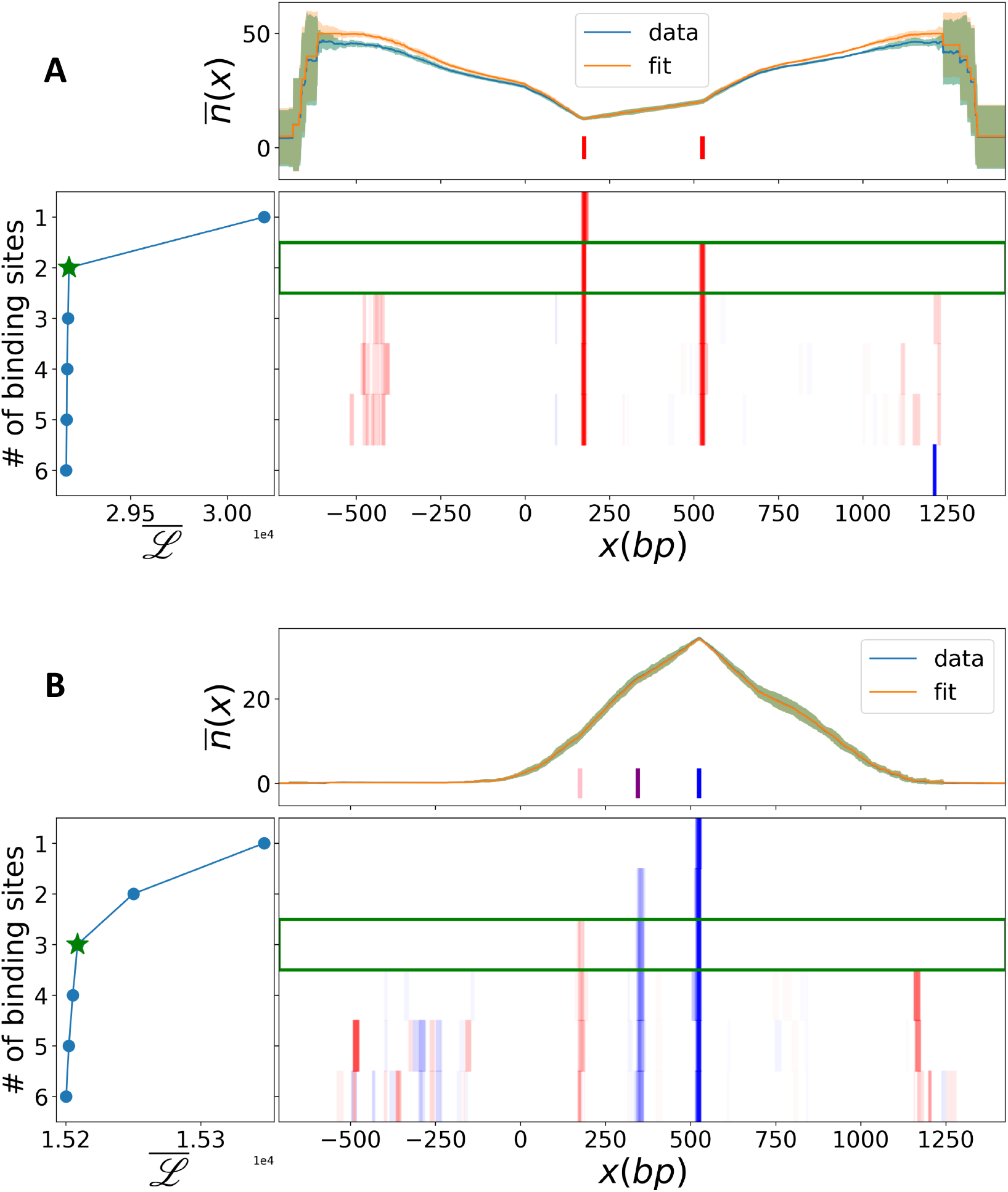
Figures A) and B) represent the dual-repressor enhancer and the inhibitor-helper-activator enhancer, respectively. Colored bars in the upper subplots show the position and width of binding sites, where red in the dual-repressor system shows the repressor site, and pink, purple, and blue in the inhibitor-helper-activator enhancer show the inhibitor, helper and activator sites, respectively. The average number of transcribed mRNA over all tiles spanning a given base pair is depicted in light blue for the data and orange for the fitted mean-field model with two binding sites in the dual-repressor enhancer and three binding sites in the inhibitor-helper-activator system, with the shade indicating the standard deviation from the mean across ten different tile configurations. The lower-left subplots depict average log-likelihood, *ℒ*, against the number of binding sites across ten configurations, with the green star marking the shoulder of the curve. In the lower-right subplots, each row shows the average weight of inferred sites at each base pair for the models with a given number of sites to infer. In these, red and blue represent positive and negative weights, respectively, indicative of a base pair’s repressive or activating role. The green rectangle shows the optimal model based on the shoulder of the *ℒ* curve. Fixed values of *r*_0_ = 50 and *N* = 400 were used to simulate both systems.

**S2:**
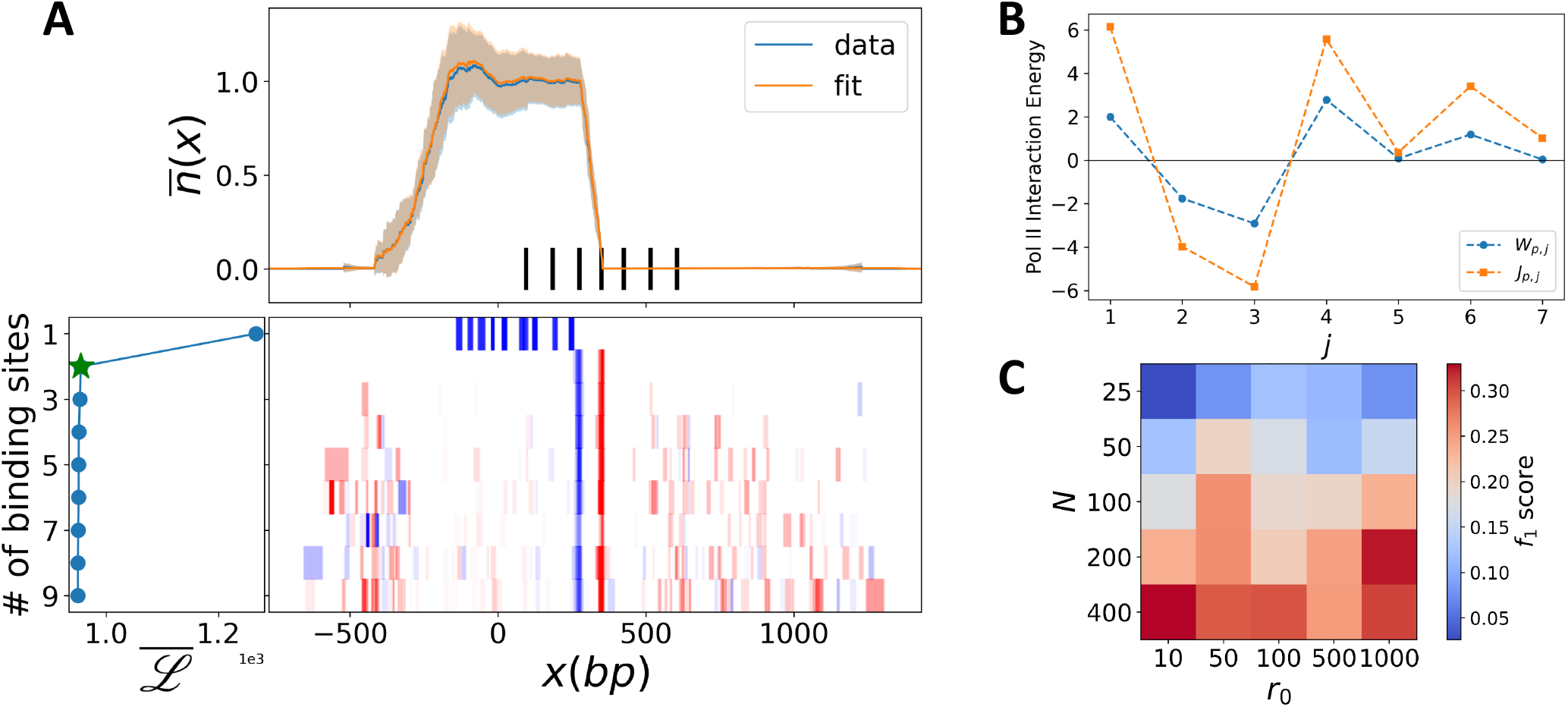
A) Black bars in the upper subplot show the position and width of binding sites (please see Supplementary Material for the exact values of locations and widths). The blue curve represents the average output over tiles of the system for the case where *N* = 400 and *r*_0_ = 50. The orange curve shows the same quantity estimated by expected count of each tile 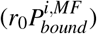 from the selected model, which here is the two-site model. In these, red and blue represent positive and negative weights, respectively, indicative of a base pair’s repressive or activating role. B) The figure shows the interaction energy of each binding site with Pol-II. *J*_*p, j*_ is the interaction energy (orange dots), while 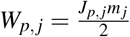 is the effective interaction energy with the *m*_*j*_ calculated from the full model (Eq. 2), but not the mean-field model. C) A heat map showing the quality of classifying the original enhancer (*f*_1_-score) as a function of the number of tiles, *N*, and output level *r*_0_.

**S3:**
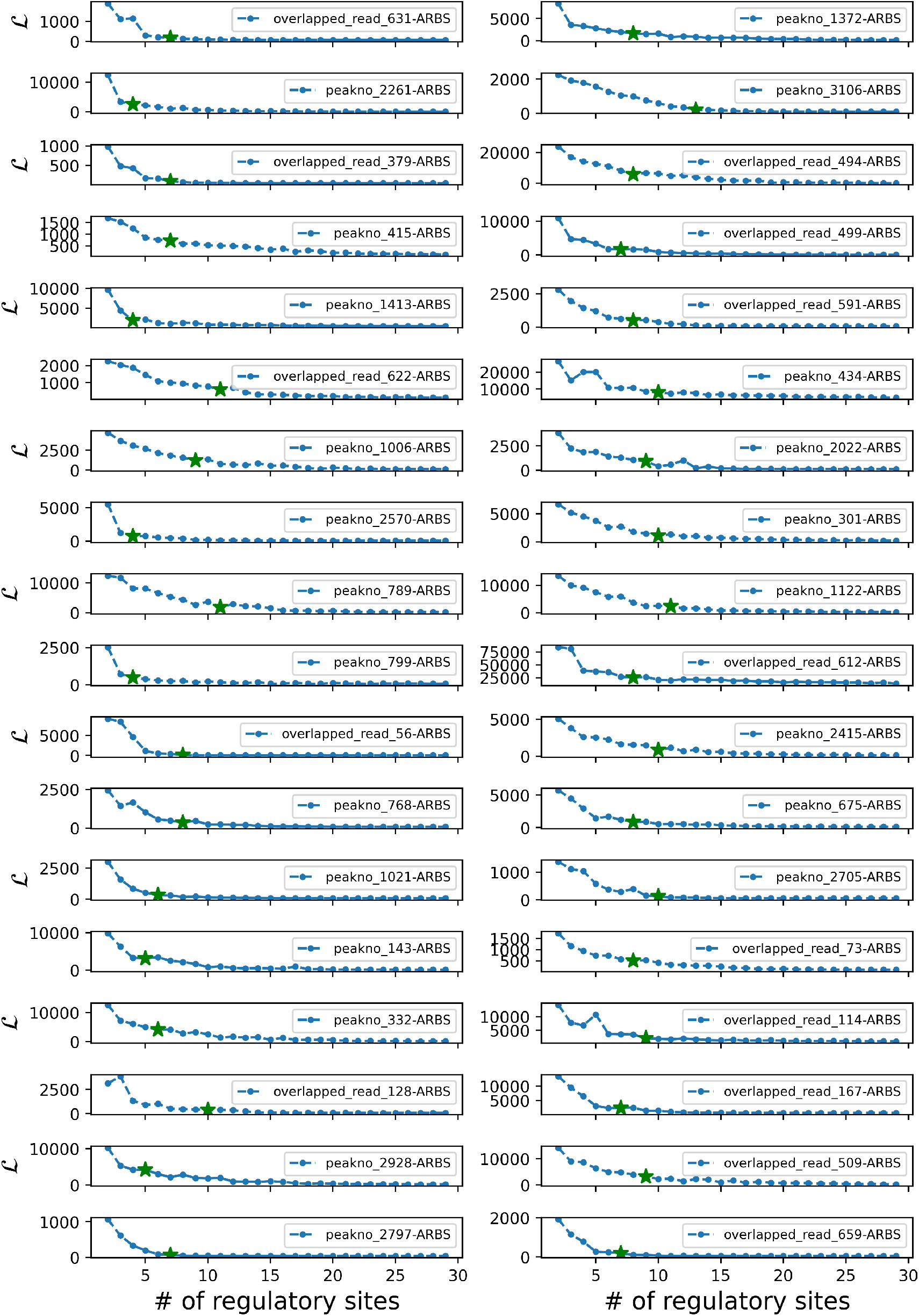
Log-likelihood versus number of regulatory sites for AR-bound regions. The green star shows the estimated location of the elbow for each region, which represents the selected model.

### 6.1 Values of used parameters in simulations

In all the following items, *h*_*i*_ shows the binding energy of the *i*-th binding site (from the left to the right of the region) in *K*_*b*_*T*, where the last component represents the promoter energy. *J*_*i j*_ shows the interaction energy between the *i*-th and the *j*-th sites. *x*_*i*_ shows the starting position of the *i*-th site. Width of all binding sites is 10 base pair in all synthetic architectours.

Length of tiles in all systems are randomly drawn from a uniform distribution with the mean, *µ*_*L*_ = 528, and the standard deviation, *σ*_*L*_ = 77. These numbers are chosen based on the tile length in the original STARR-seq assay.

- Helper-activator

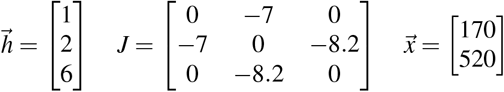
- Inhibitor-repressor

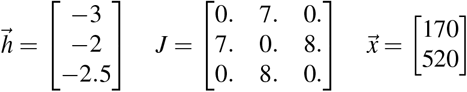
- Dual repressor

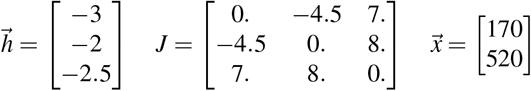
- Inhibitor-helper-activator

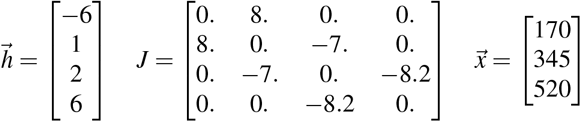
- Complex enhancer 1 (Fig. 4)

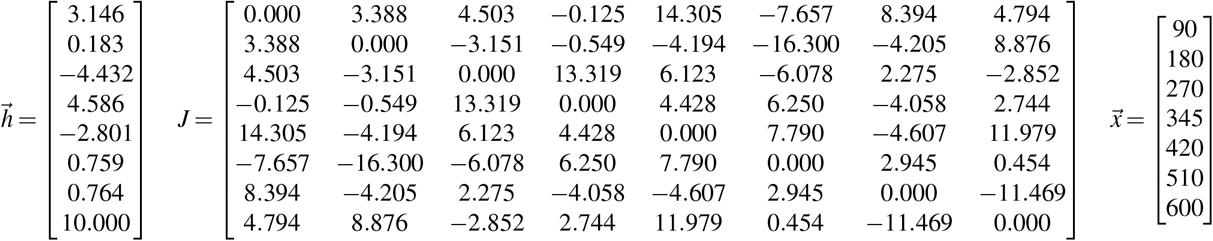
- Complex enhancer 2 (Fig. S2)

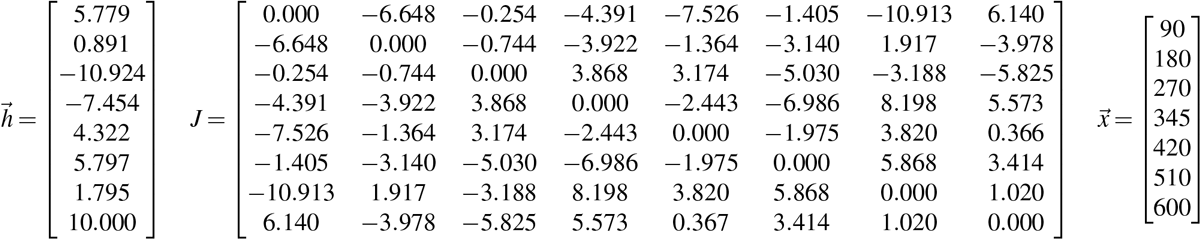
- AR-bound regions

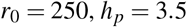

